# Variation of bacterial communities along the vertical gradient in Lake Issyk Kul, Kyrgyzstan

**DOI:** 10.1101/864355

**Authors:** Keilor Rojas-Jimenez, Alex Araya-Lobo, Fabio Quesada-Perez, Johana Akerman-Sanchez, Brayan Delgado-Duran, Lars Ganzert, Peter O. Zavialov, Salmor Alymkulov, Georgiy Kirillin, Hans Peter Grossart

## Abstract

In this study, we explored the diversity and community composition of bacteria along a vertical gradient in Lake Issyk Kul, Kyrgyzstan, one of the largest and deepest brackish lakes in in the world. We identified 4904 bacterial ASVs based on analysis of 16S rRNA gene sequences and determined significant changes in the composition responding mainly to the variables depth and salinity. A higher abundance of Proteobacteria and Bacteroidetes was observed in the surface waters and the lake tributaries. Cyanobacteria were more abundant in the deep chlorophyll maximum (28.5 to 128 m), while Planctomycetes and Chloroflexi were dominant at depths between 128 to 600 m. According to our machine learning analysis used for identifying the most critical environmental factors, depth and temperature revealed the strongest effect on members of Proteobacteria, Planctomycetes, and Chloroflexi, while oxygen is associated with the variations in Cyanobacteria. Also, a notable increase in alpha diversity estimations was observed with increasing water depth. This work evidences significant differences in the structure of bacterial communities along the depth gradient in deep, transparent lake ecosystems. Notably, there is a dominance of Planctomycetes and Chloroflexi in the deepest layers, which can only be seen in a few other lakes with similar characteristics as Lake Issyk Kul and raises questions about their ecological role.

## Introduction

The current knowledge about the distribution of planktonic bacterial communities along the vertical gradient in lake ecosystems is limited, particularly in lakes with depths greater than 200 m. Of the total existing lakes globally, only a few contain mesopelagic ecosystems (200 to 1000 m). Of these, only a handful has been studied concerning diversity and composition of their microbial communities, with examples mainly from Lake Baikal in Russia (Glöckner et al. 2000; Bel’kova et al. 2003; Kurilkina et al. 2016), Lake Tanganyika in Africa (De Wever et al. 2005), and Crater Lake in the United States (Urbach et al. 2001).

Water depth is one of the main factors that influences the composition and distribution of bacterial communities in both oceans and lakes. Along the vertical gradient, there is a transition from the photic to the aphotic zone, which also includes a decrease in temperature, oxygen and nutrients, concomitant with an increase in hydrostatic pressure (DeLong et al. 2006; Bryant et al. 2012; Garcia et al. 2013; Mende et al. 2017; Wu et al. 2019). Salinity is another factor that influences the composition of microbial communities in aquatic ecosystems in both the vertical and horizontal gradients (Herlemann et al. 2011; Newton et al. 2011; Rojas-Jimenez et al. 2019).

Recent applications of high-throughput sequencing technologies are providing a more in-depth insight into the taxonomic structure and spatial distribution of bacterial communities, offering new opportunities to analyze the complexity of microbial communities in aquatic ecosystems (Shaw et al. 2008; Van Der Heijden et al. 2008; Caporaso et al. 2011; Newton et al. 2011). In addition to the advances in sequencing technologies, Machine Learning tools contribute to reveal unexpected biological patterns, discover ecological relationships between microorganisms and their environment, as well as in formulating new biological hypotheses (Eraslan et al. 2019; Qu et al. 2019).

In this work, we studied the spatial variability and distribution of the bacteria communities of Lake Issyk Kul. This lake is located in the northeast of Kyrgyzstan and has the singularity of being one of the largest and deepest lakes in Central Asia and even worldwide. It is a mountain lake with an average altitude of 1,609 meters above sea level, and with a maximum depth of 702 m. Lake Issyk Kul occupies a closed basin of tectonic origin with a total area of 6,280 km^2^ and is considered the second largest mountain lake in the world after Lake Titicaca in Bolivia. Lake Issyk Kul arouses great scientific interest, in particular, because of its geophysical and ecological characteristics. Being the six-deepest lake of the world and tenth-largest by volume, it reveals surprisingly high rates (below 11 years) of deep-water renewal (Hofer et al. 2002; Peeters et al. 2003), which maintain the lake water column well-oxygenated down to its deepest parts (Zavialov et al. 2018). Despite the absence of outflows, the salinity of the lake water of 6 g kg^-1^ is rather low, which suggests the lake to become endorheic relatively recently, possibly due to tectonic processes (Romanovsky 2002). Lake Issyk Kul is (ultra)-oligotrophic with highly transparent water, reaching Secchi depths of >20 m. The low nutrients content in the surface waters and the high amount of solar radiation penetrating deep into the water column creates a unique environment, characterized by a maximum of photosynthesis at depths of 30-40 m. First systematic studies on Lake Issyk Kul date back to the 1920s. It was one of the most studied lakes in the 1960-80s, but research stopped abruptly after the collapse of the Soviet Union. During the 1960-80s the lake ecosystem experienced a substantial impact by human activities, including the introduction of non-native commercial fish species displacing the endemic species, input of wastewater and pollution by industry and agriculture, with effects on the aquatic ecosystem that remain-unknown (Savvaitova and Petr 1992; Giralt et al. 2004; Baetov 2005).Yet, almost nothing is known about microbial communities, in particular heterotrophic bacteria at the base of the food web.

Therefore, we used Illumina sequencing of the 16S rRNA gene to characterize the bacterial populations inhabiting Lake Issyk Kul and some of its tributaries. We also analyzed the effect of different environmental variables on the structure of bacterial communities, using various statistical tools and machine learning techniques. We hypothesized that the stratification of the water column will affect the composition of bacteria, where the deeper layers will be characterized by poorly described taxa with their ecological roles being little known. Additionally, this work will allow us to compare the composition of the bacterial community in the rivers with respect to that of the lake.

## Materials and Methods

### Environmental variables and sampling

The lake expedition was carried out during the cruise of the RV Moltur from June 25 to July 1, 2017. All sampling stations in the lake and its tributaries are shown in **Figure 1**. Sampling and measurement of physical and chemical variables was performed as previously described (Zavialov et al. 2018). Briefly, water samples were taken using HydroBios 5 L Niskin bottles at all stations at different depth levels from the surface to the bottom. Depth was measured using a digital sounder (model LMS-350, Lowrance). Profiles of temperature, electric conductivity, and fluorescence were taken with the CTD-probe SeaBird 19plus (Sea-Bird Scientific, USA). Oxygen profiles were taken with the fast-response oxygen optode Rinko-I (JFE Advantech, Japan). Salinity was calculated from temperature, electrical conductivity, and pressure using the empirical formula for the Lake Issyk Kul ionic composition proposed by Peeters et al. (2003). Chemical determinations were performed in the laboratory using standard techniques; Winkler method for oxygen; potentiometric method for pH; titration method for alkalinity; colorimetric assay method for phosphates, silicates, nitrates, nitrites, and carbon (CO_2_, CO_3_, pCO_2_) in different carbonate equilibria (**Suppl. Fig. 1**).

**Figure 1.**
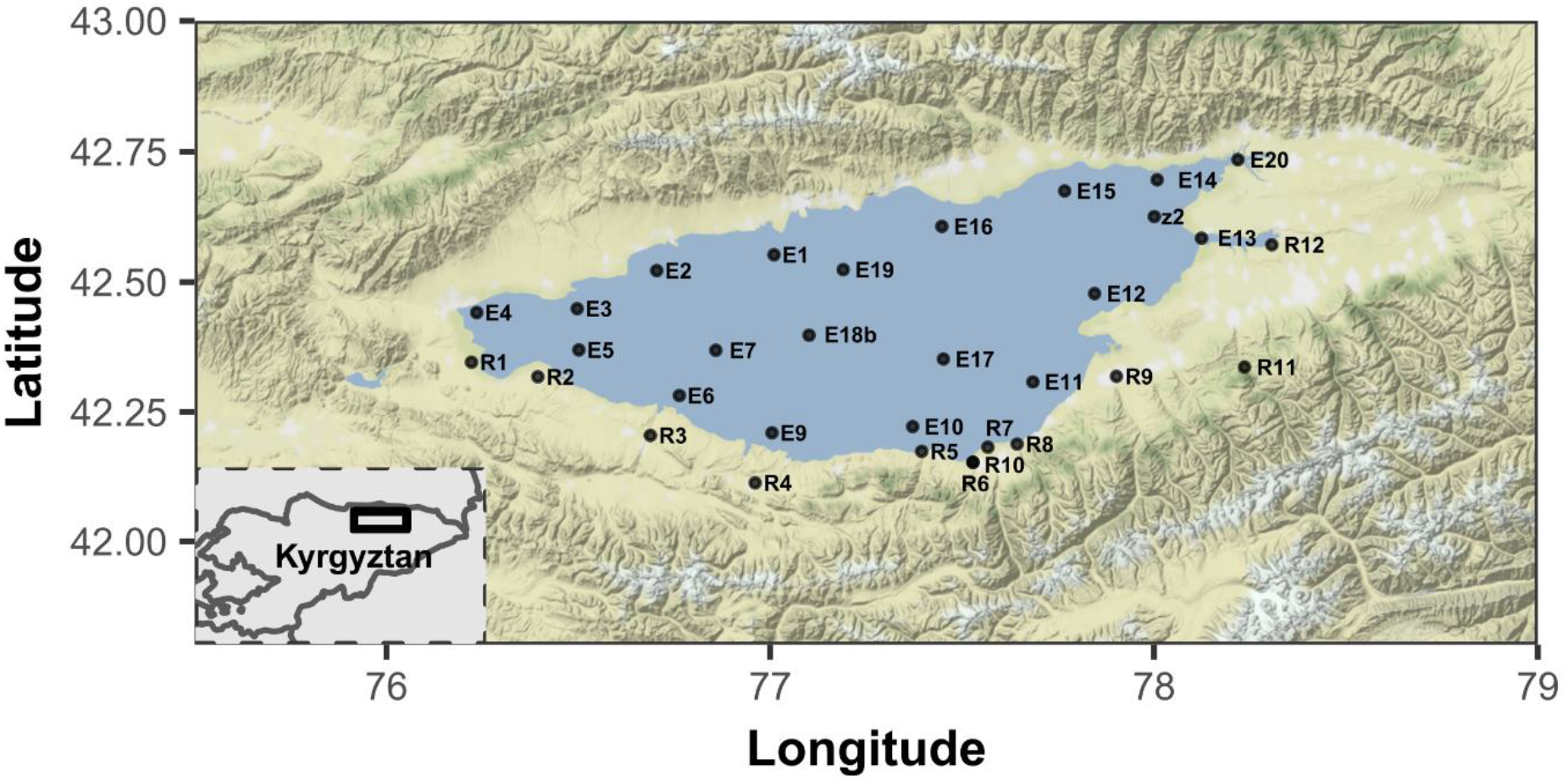
Location of sampling points in Lake Issyk Kul, Kyrgyztan and some of its tributary rivers.

### Molecular analyses

Samples were immediately filtered on board after retrieval of the Niskin bottle. The water samples were filtered through 0.22 um Sterivex® filters (EMD Millipore, Germany) connected to a peristaltic pump (EMD Millipore, Germany) to concentrate bacteria and subsequently stored at −20 °C. DNA was extracted from 0.22 μm Sterivex filters using the QIAamp DNA mini kit (Qiagen, Germany) following the protocol for tissue with some modifications. Briefly, the filters were cut into pieces and put into a 2 mL tube. A mix of zirconium beads and 360 μL of buffer ATL was added and vortexed for 5 min at 3,000 rpm in an Eppendorf MixMate^®^ (Eppendorf, Germany). Proteinase K (>600 mAU/ml, 40 μL) was added and incubated at 57 °C for 1 h. After centrifugation for 1 min at 11,000 rpm, the supernatant was transferred to a new 2 mL tube, and extraction was performed following the manufacturer’s protocol. PCR, library preparation, and sequencing were done by LGC Genomics (Berlin, Germany). Briefly, the V3-V4 region was amplified using primers 515F-Y/926R (Parada et al. 2016) followed by library preparation (2 × 300 bp) and sequencing on a MiSeq Illumina platform. Sequences were quality checked and analyzed using version 1.12 of the DADA2 pipeline (Callahan et al. 2016). This process resulted in an amplicon sequence variant (ASV) table, a higher-resolution analogue of the traditional OTU table, which records the number of times each exact amplicon sequence variant was observed in a sample. Taxonomy assignment was performed by comparing sequences against the SILVA reference database v132 (Quast et al. 2013) and then curated by comparing sequences against NCBI. Global singletons were removed. Samples with a low sequencing depth (< 2000 sequences) were also discarded from the analysis. After these processes, we obtained 1,790,675 sequences and 4904 bacterial ASVs. The average number of sequences per sample was 24470 ranging from 3435 to 82992. The sequence data were deposited in GenBank (https://www.ncbi.nlm.nih.gov/genbank/) under the SRA accession numbers xxx.

### Statistical analysis

The statistical analyses and their visualization were performed in the R Project statistical program (R-Core-Team 2019), using the Rstudio interface. The dlookr package was used to transform the numerical variables depth (*binning* function adjusted with kmeans algorithm) and salinity (*binning* function: the categorization of this variable was carried out observing the threshold at which the abundance of the bacterial species changed considerably) into categorical variables. The Vegan package (Oksanen et al. 2017) was used for non-metric multidimensional scaling (NMDS) and permutational analysis of variance (PERMANOVA; *adonis2* function with 999 permutations with Bray-Curtis adjustment). To adjust the pairwise p-values, the *pairwise.adonis* function with Bonferroni adjustment was used. The NMDS was represented in a two-dimensional plot based on a Bray-Curtis similarity matrix.

We applied random forest models to identify the critical environmental parameters affecting the relative abundance of the seven most abundant bacterial phyla utilizing package randomForest (Liaw and Wiener 2002). The degree of importance of each predictor variable was determined by quantifying the effect of its removal on the model’s accuracy across all trees (number of trees=10001). To assess model significance, tests with 1000 permutations were employed. The relationships between environmental variables and bacterial family richness as well as diversity (represented by the Shannon Index) were explored with generalized additive models (GAM), using the mgcv and mgcViz packages (Wood and Wood 2015; Fasiolo et al. 2019), and with Kruskal-Wallis tests by rank and Mann-Whitney tests for comparisons between the variables. Finally, random forest models were applied to identify the critical environmental factors affecting the richness and diversity of the community (number of trees=10001).

## Results

We observed pronounced changes in the composition of the bacterial communities of Lake Issyk Kul along the vertical gradient but not the horizontal gradient, responding mainly to the variables depth and salinity (**Figure 2**). In total, 4904 bacterial ASVs were identified, where the phylum Proteobacteria was the most abundant, representing in average 28.49% of all sequences. In lower average proportions, other groups were also detected, such as Bacteroidetes (16.76%), Cyanobacteria (14.81%), Planctomycetes (12.85%), Actinobacteria (9.82%), Verrucomicrobia (8.67%), and Chloroflexi (6.56%) (**Suppl. Fig. 2**). The primers set used also allowed the identification of Archaea, particularly of the genus *Nitrosopumilus* (Thaumarchaeota), but in a very low proportion with respect to bacteria.

**Figure 2.**
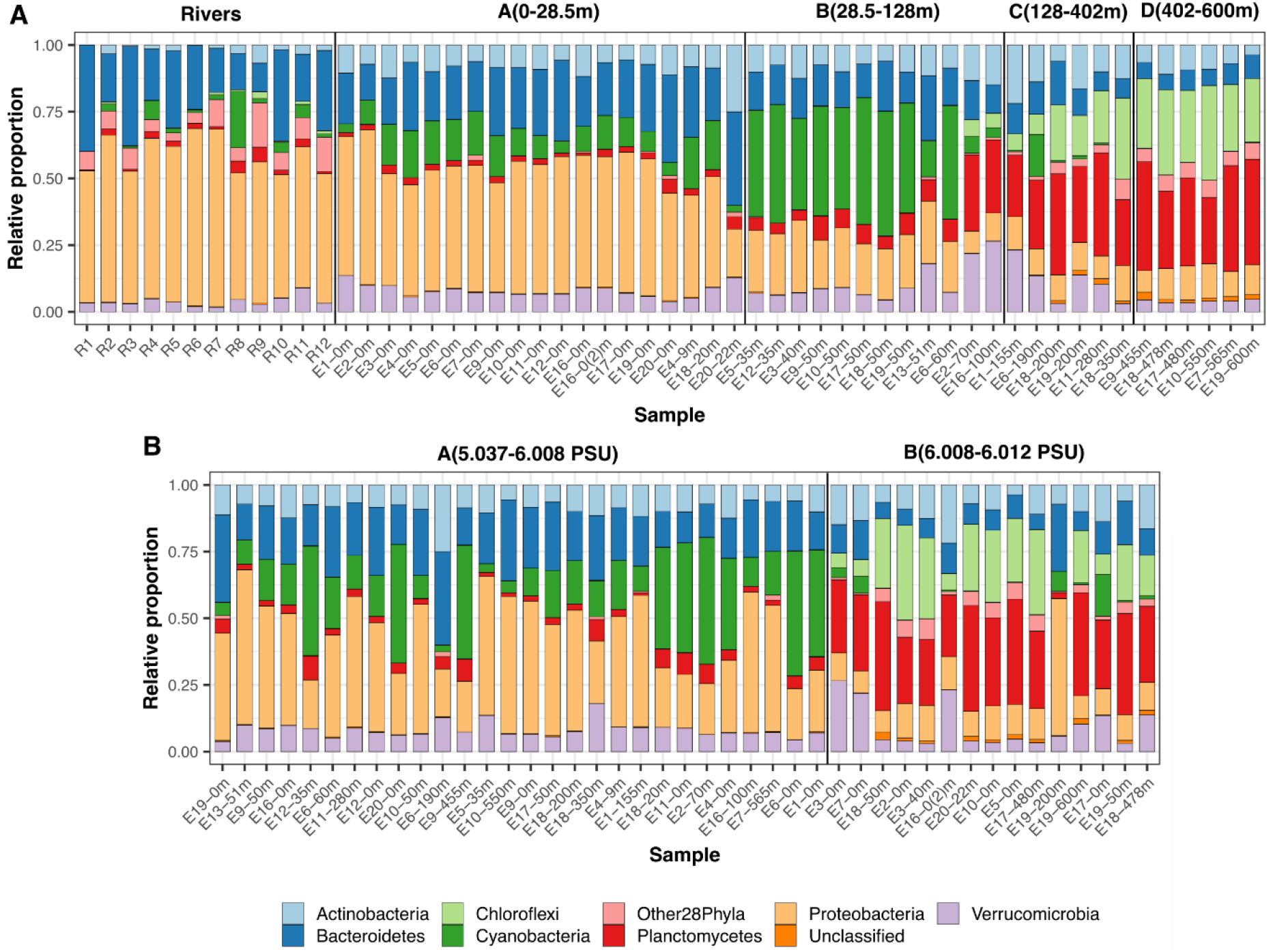
Composition of bacterial communities, determined at the taxonomic level of the phylum, for several water samples from Lake Issyk Kul, Kyrgyzstan. Composition is shown according to depth (**A**) and salinity (**B**) gradients.

We used the K-means algorithm to create clusters of samples according to depths, distinguishing four layers: 0 to 28.5 m, 28.5 to 128 m, 128 to 402 m, 402 to 600 m. Another cluster included the tributary rivers. This grouping was consistent with differences observed in their bacterial composition (**Figure 2A**). At the phylum level, a greater abundance of Proteobacteria and Bacteroidetes was found in the surface waters of the lake and the river samples. In the deep chlorophyll maximum (DCM, between 28.5 to 128 m), there was a higher abundance of Cyanobacteria, while Planctomycetes and Chloroflexi became more abundant at greater depth, i.e. between 128 to 600 m.

At the ASV level, a strain associated with the genus *Sphingomonas* (Alphaproteobacteria) was the most abundant in the rivers. *Cyanobium* (Cyanobacteria), *Loktanella* (Alphaproteobacteria), and an unknown strain of Burkholderiaceae (Gammaproteobacteria) were the most abundant in the surface waters of the lake. *Cyanobium* also dominated in the DCM between 28.5 to 128m. Three strains related to families Anaerolineaceae (Chloroflexi) as well as Phycisphaeraceae and Gimesiaceae (Planctomycetes) were the most abundant ASVs in the deeper layers (128 to 402 m and 402 to 600 m) although with small differences between each layer.

The observed variation in the composition of bacterial communities along the depth gradient was consistent with the NMDS analysis that showed a clear separation of the clusters (**Figure 3**). These results were also compatible with the Permanova analysis, showing statistically significant differences between the bacterial communities of the rivers and the lake, and even within each of the four depth clusters identified (**Suppl. Table 1**).

**Figure 3.**
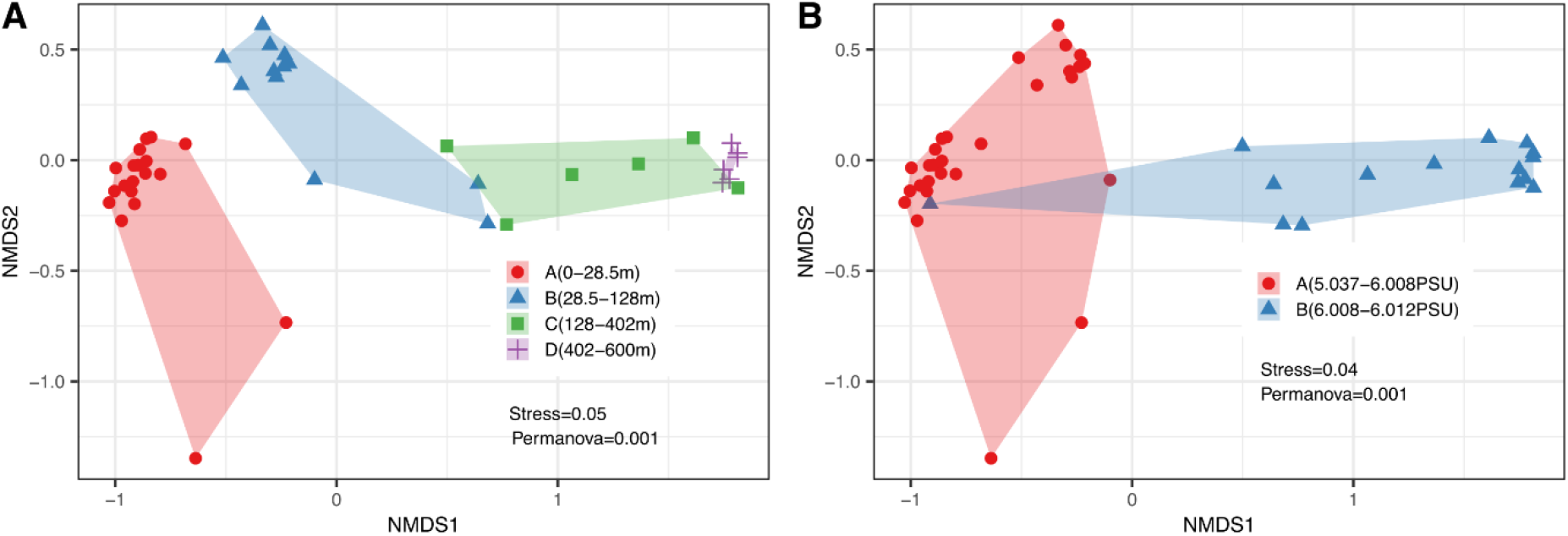
Non-metric multidimensional scaling analyses of the bacterial communities in Lake Issyk Kul performed by depth (A) and salinity (B).

We also observed differences in the composition of bacterial communities, according to salinity (**Figure 2B**). In superficial layers, salinity ranged between 5.037 and 6.008 PSU, where a higher abundance of Proteobacteria, Cyanobacteria, and Bacteroidetes was observed. The river samples were not included in this analysis because the information of this variable was not recorded. In the deeper layers, salinity was more stable, between 6.008 and 6.012 PSU, where a higher relative abundance of Planctomycetes and Chloroflexi was observed. These differences in community composition were consistent with the clustering of the NMDS analysis (**Figure 3B**), and the Permanova analysis, showing significant differences in the composition between these two salinity layers (**Suppl. Table 1**). Regarding other environmental parameters (such as oxygen or temperature), we did not find any significant effect on the composition of the lake bacterial communities.

As a complementary method, we used machine learning (Random Forest algorithm) to identify environmental variables that have more weight in the observed variation of the relative abundance of the bacterial phyla (**Figure 4**). According to this analysis, depth and temperature are the environmental factors that have the most substantial effect on the variations in Proteobacteria, Planctomycetes, and Chloroflexi. The higher abundance of Cyanobacteria was associated with conditions of high oxygen, while for Actinobacteria and Verrucomicrobia no specific variables were distinguished as responsible for the variations in their abundance.

**Figure 4.**
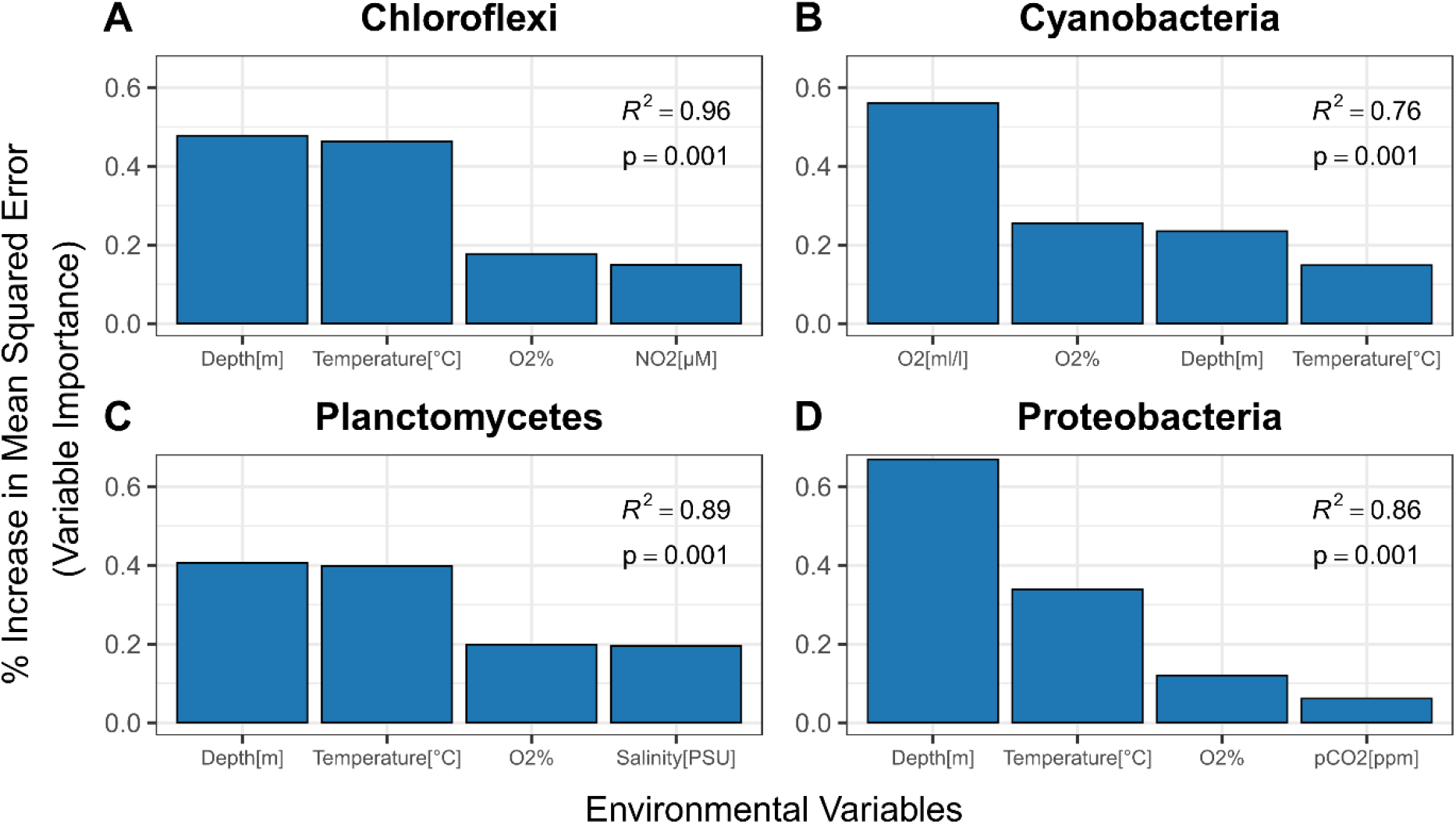
Determination of the influence of environmental variables on the variations in abundance of different bacterial groups. The analyses were performed using Random Forest, considering the most abundant bacterial groups per depth layer, including Chloroflexi (**A**), Cyanobacteria (**B**), Planctomycetes (**C**), and Proteobacteria (**D**).

We also used machine learning and generalized additive models (GAM) to analyze two alpha diversity estimators: richness and Shannon Index. The richness, estimated at the taxonomic level of family, ranged between 75 and 342 per sample. The factor that best explained this variability was depth (Random Forest; p = 0.001, **Figure 5A**). We observed an increase in richness with increasing depth, ranging from 110 ± 7 bacterial families in surface waters to 309 ± 6 in the deepest layers (**Figure 5B**). The variation in richness between layers was statistically significant (Kruskal-Wallis, p <0.001), but according to the GAM, there was a stabilization in the number of families at 280 m (GAM; R2 = 0.827, F = 41.89, p <0.001, **Figure 5C**). At lower temperatures, consistent with greater depths, we also observed an increase in richness (**Figure 5D and Figure 5E**).

**Figure 5.**
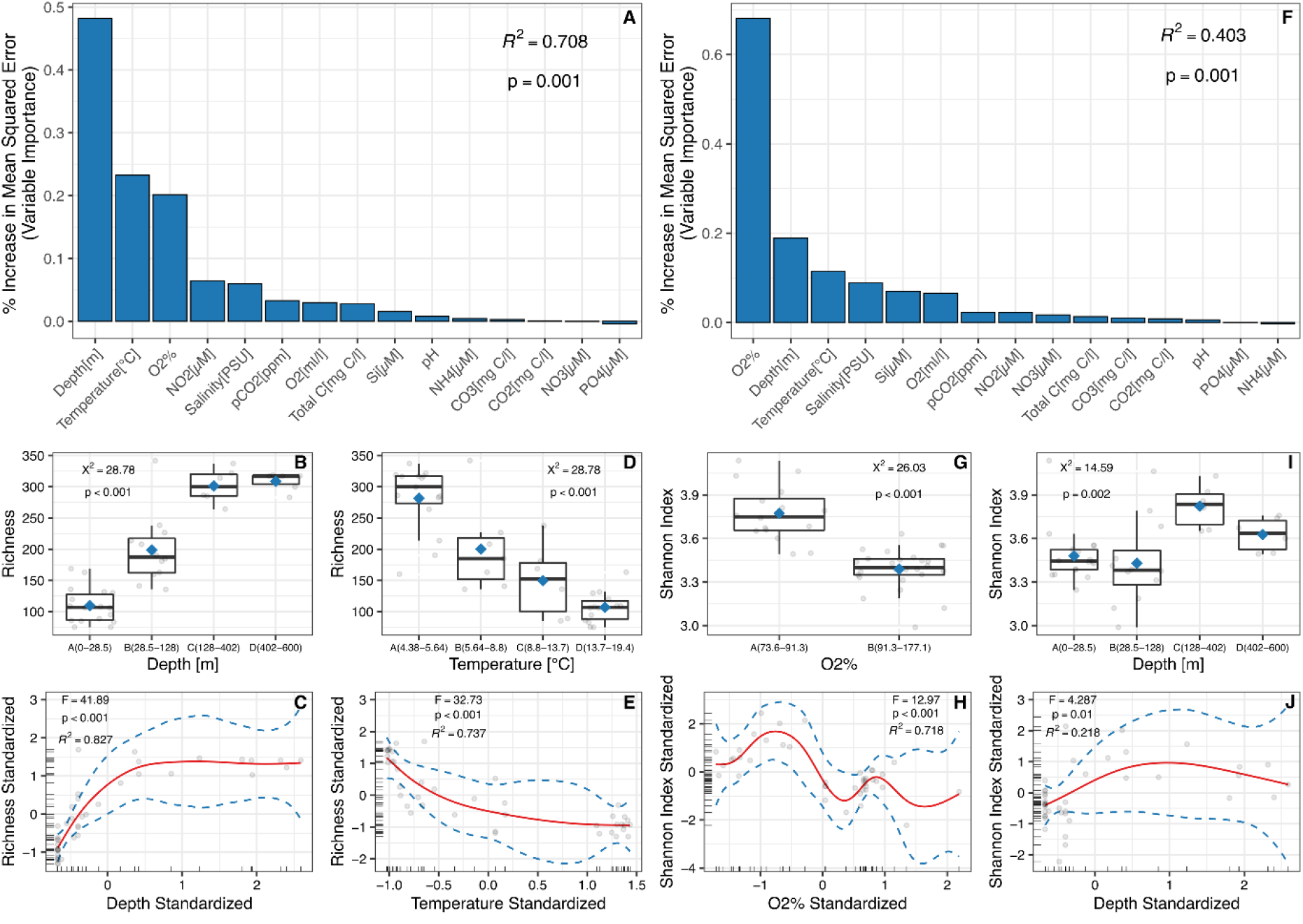
Determination of the influence of environmental variables on the variations in richness at the family level (**A**) and Shannon Index (**J**), determined with Random Forest. The boxplots show the alpha diversity values according to the main environmental variables (**B, D, G, I**) with standardized values considering the GAM model predicted relationships (**C, E, H, J**).

The percentage of oxygen (O_2_ %) was the factor that explained best the variability in the values of the Shannon Index among all measured environmental factors (Random Forest; p = 0.001, **Figure 5F**). Although Lake Issyk Kul presents aerobic conditions along the vertical gradient (**Supplementary Figure 3**), higher diversity values were determined in the layers where oxygen was lower (73.6-91.3%), while diversity values were lower in the superficial and DCM layers, where the oxygen is higher (91.3-177.1%). Therefore, we observed a slight decrease in diversity as the oxygen level increases (GAM; R2 = 0.718, F = 12.97, p <0.001, **Figure 5G and Figure 5H**). This variation in the Shannon Index between oxygen levels was statistically significant (Kruskal-Wallis; χ^2^ = 23.61, p <0.001). Regarding the variation of this index concerning depth, we observed that the highest values of diversity were found between 128-402 m (**Figure 5I, Figure 5J**).

## Discussion

This is the first study that describes the diversity and composition of bacterial communities in Lake Issyk Kul, Kyrgyzstan, using next generation sequencing and a high number of sampling points along the vertical gradient. The main finding is that the depth plays a decisive role in the variations of the community structure, which is consistent with the results of other studies conducted in deep lakes such as Lake Baikal in Russia, Crater Lake in the United States, and Lake Tanganyika in Central Africa (Glöckner et al. 2000; Urbach et al. 2001; Bel’kova et al. 2003; De Wever et al. 2005; Kurilkina et al. 2016) . However, Lake Issyk Kul showed some differences in terms of the composition of the most abundant groups in the different layers, probably related to particular brackish conditions and high oxygenation.

The composition of the bacterial communities in the tributaries was significantly different from that found in the different layers of the lake, including the surface layer. For example,*Sphingomonas* was the most abundant genus in the rivers while *Loktanella* and an unknown Burkholderiaceae were the most abundant in the surface layer of the lake. Also, Cyanobacteria, Planctomycetes, and Verrucomicrobia were almost missing in the rivers, possibly due to flowing water conditions and high sediment and nutrients load. Although this work was not designed to determine the role of rivers in the provision of seeding communities, nor to assess the effects of human activities on the lake, the results showed clear differences between these two ecosystems. It will be important, however, to continue monitoring the impact of potential sources of pollution, such as the cities located in the northern sector of Lake Issyk Kul.

With the increase in depth and slightly more saline conditions, we observed a decrease in the abundance of Proteobacteria, which is consistent with previous studies showing the effects of these environmental variables on the abundance of this phylum (Holdridge 1960; Simon et al. 1999; Cottrell and Kirchman 2003; Kirchman et al. 2005; Kan et al. 2008). The DCM was dominated by Cyanobacteria and particularly by the genus *Cyanobium*, which is consistent with previous observations in other oligotrophic lakes in Europe (Padisák et al. 2003, 2004; Selmeczy et al. 2016). Such a deep DCM could be the result of extremely high subsurface radiation and low nutrients, which is also concurrent with maximum values of oxygen and chlorophyll a.

Previous studies have shown the presence of Chloroflexi and Planctomycetes in deep waters (Urbach et al. 2001; De Wever et al. 2005; Breuker et al. 2013; Campbell and Kirchman 2013; Luo et al. 2014; Kurilkina et al. 2016; Li et al. 2018), however, this is the first study showing the dominance of these two phyla, and more specifically of Anaerolineaceae and Phycisphaeraceae-Gimesiaceae, in the deeper layers of a lake. The microorganisms in the deep waters of Lake Issyk Kul face aphotic conditions, higher salinity than in the surface, lower temperatures, high hydrostatic pressure, and high availability of oxygen. Therefore, it is possible that the bacteria inhabiting this ecosystem should have some physiological adaptations to tolerate higher hydrostatic pressures and higher salinity.

The hydrostatic pressure at 600 m could trigger some mechanisms for adaptation such as the increase in the proportion of unsaturated fatty acids in the phospholipids of membranes (Delong and Yayanos 1985; DeLong and Yayanos 1986; Tamburini et al. 2013; Wannicke et al. 2015), systems for active transport of sugars (DeLong and Yayanos 1987), and increase in bacterial protein production (Somero 1992; Wannicke et al. 2015). It is also possible that physiological adaptations to depth could be accompanied by changes at the genomic level, including larger genome size, higher genomic GC content, and proteins with higher nitrogen but lower carbon content (Mende et al. 2017). Proposing the occurrence of these adaptation mechanisms in microorganisms of the deep waters of Lake Issyk Kul would be somewhat theoretical; however, it deserves further studies given that the highest richness and diversity was found in the deepest part of the ecosystem. For this, it will be important to include novel culture techniques, microscopy, physiological tests, and genomic analysis.

In conclusion, in this work we demonstrate the effect of depth and salinity on the variations of the structure of bacterial communities along the vertical gradient in Lake Issyk Kul, which is consistent with previous studies (Koizumi et al. 2003; Bryant et al. 2012; Liu et al. 2014; Mestre et al. 2017; Wu et al. 2019). We highlight a high bacterial diversity in the deepest layers of the lake, where microorganisms live in brackish waters, aphotic conditions, low temperatures, high hydrostatic pressure, but completely aerobic conditions. The dominance of Chloroflexi and Planctomycetes in the deeper layers raises questions about their ecological and functional roles, which should be further explored.

## Acknowledgements

We are deeply thankful to the crew of the RV Moltur and the scientific team of the Shirshov Institute of Oceanology, Moscow for assistance with fieldwork. The field work was supported by Russian Foundation for Basic Research and Russian Geographic Society through their joint Grant Nr. 17-05-41043. GK and HPG were supported by the German Research Foundation (DFG Projects Ki 853-11/2, Ki 853-13/1 and GR1540/29-1).

## Supplementary Materials

**Supplementary Table 1.** Results of the statistical analysis where the structure of the microbial communities is compared between depth layers and salinity levels. The comparison was performed using Permanova. Asterisks indicate significant differences.

| Variable | Pairs | F.Model | R<sup>2</sup> | p.value | p.adjusted |
| --- | --- | --- | --- | --- | --- |
| Depth* | A(0-28.5m) vs B(28.5-128m)* | 7.74 | 0.21 | 0.001 | 0.006 |
|  | A(0-28.5m) vs C(128-402m)* | 15.07 | 0.39 | 0.001 | 0.006 |
|  | A(0-28.5m) vs D(402-600m)* | 23.55 | 0.50 | 0.001 | 0.006 |
|  | B(28.5-128m) vs C(128-402m)* | 8.79 | 0.35 | 0.001 | 0.006 |
|  | B(28.5-128m) vs D(402-600m)* | 18.72 | 0.53 | 0.001 | 0.006 |
|  | C(128-402m) vs D(402-600m)* | 4.14 | 0.29 | 0.029 | 0.174 |
|  | A(0-28.5m) vs Rivers* | 32.31 | 0.52 | 0.001 | 0.01 |
|  | B(28.5-128m) vs Rivers* | 21.53 | 0.49 | 0.001 | 0.01 |
|  | C(128-402m) vs Rivers* | 11.86 | 0.42 | 0.001 | 0.01 |
|  | D(402-600m) vs Rivers* | 14.89 | 0.48 | 0.001 | 0.01 |
| PSU* | A(3.979-4.746) vs B(4.746-4.749)* | 22.50 | 0.35 | 0.001 | 0.001 |

**Supplementary Figure 1.**
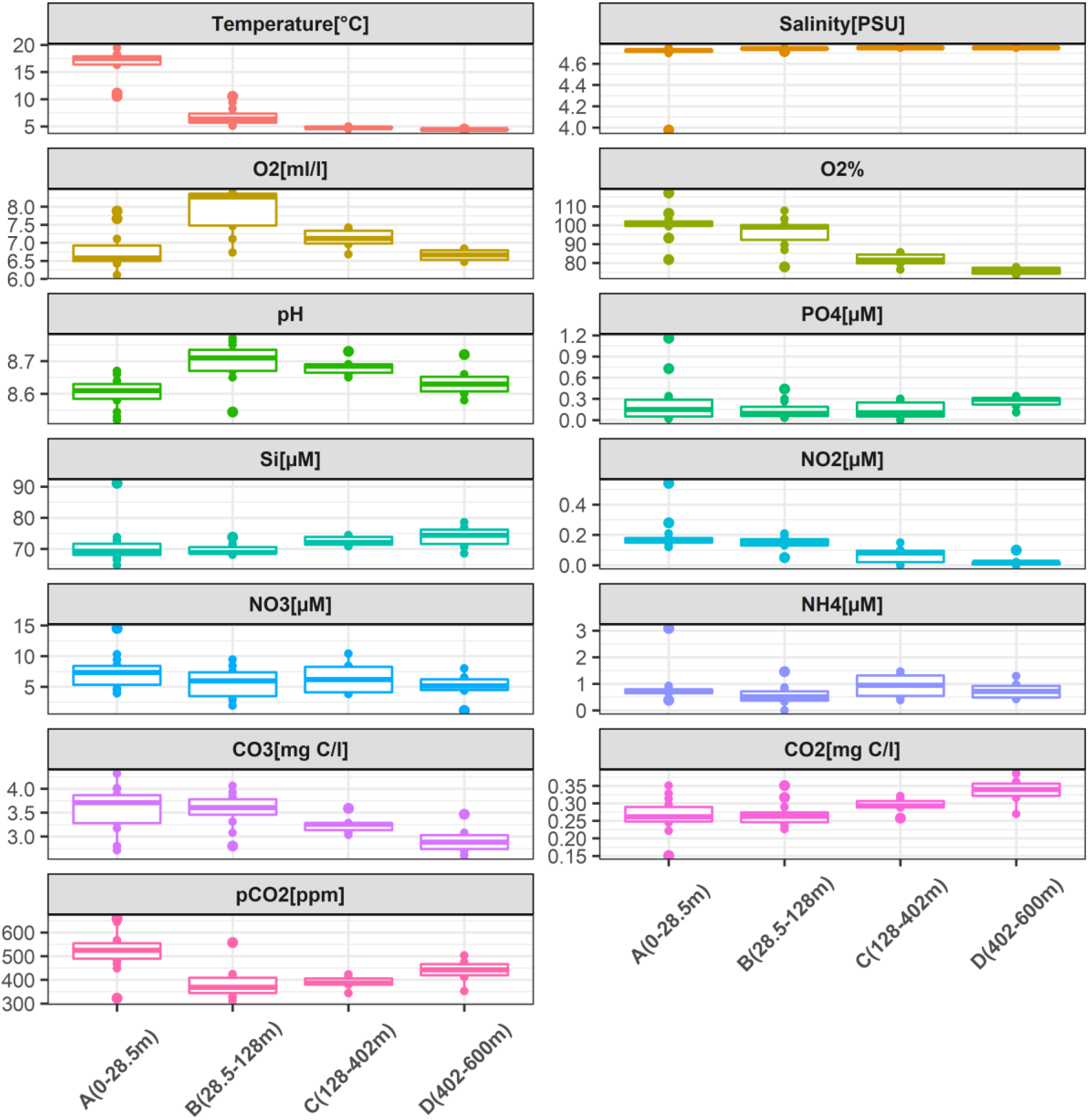
Box plots of the measured environmental variables at different points of Lake Issyk Kul, by depth layers.

**Supplementary Figure 2.**
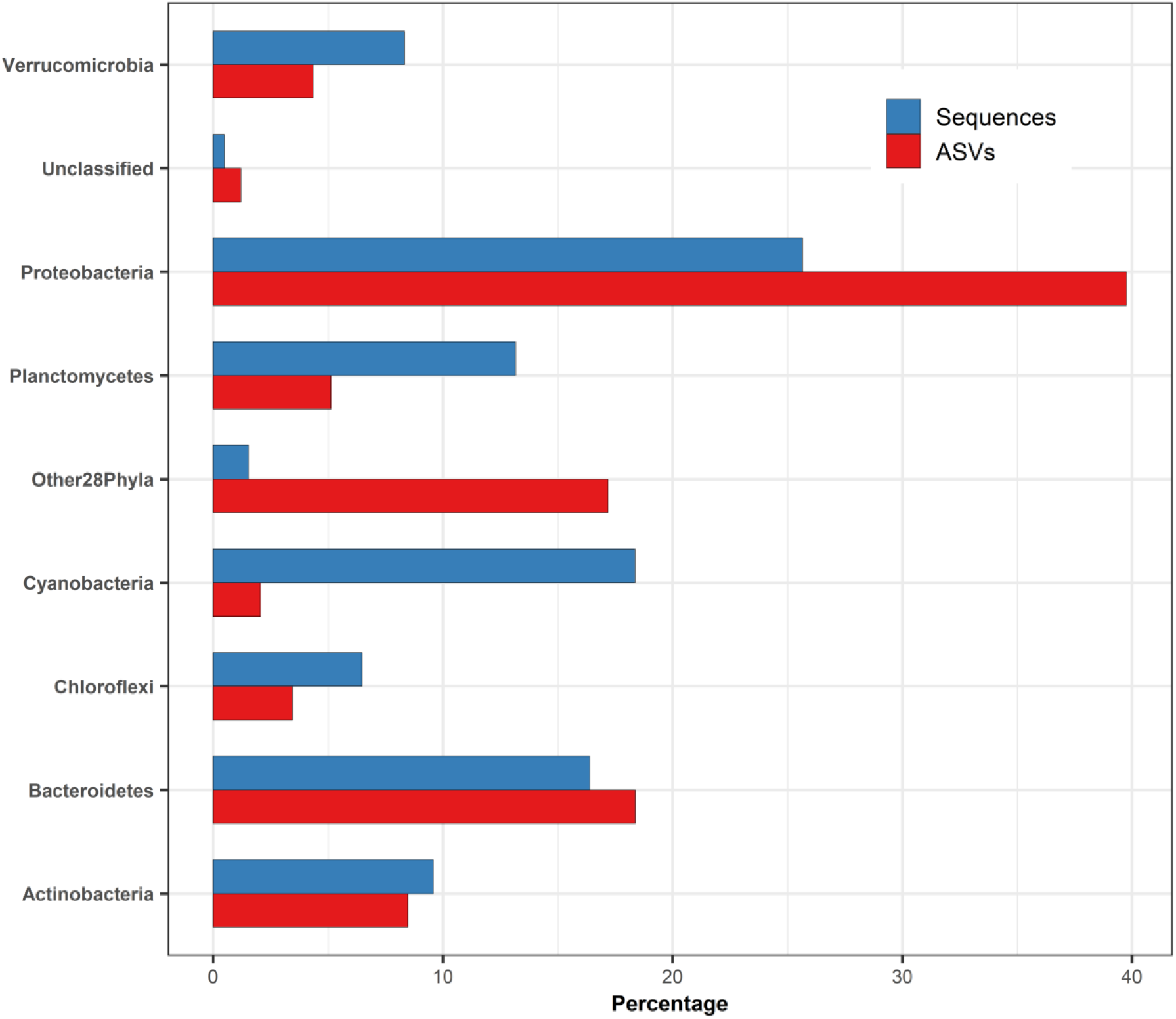
Relative abundance of the main bacterial groups in Lake Issyk Kul, shown in terms of the number of ASVs and number of sequences.

**Supplementary Figure 3.**
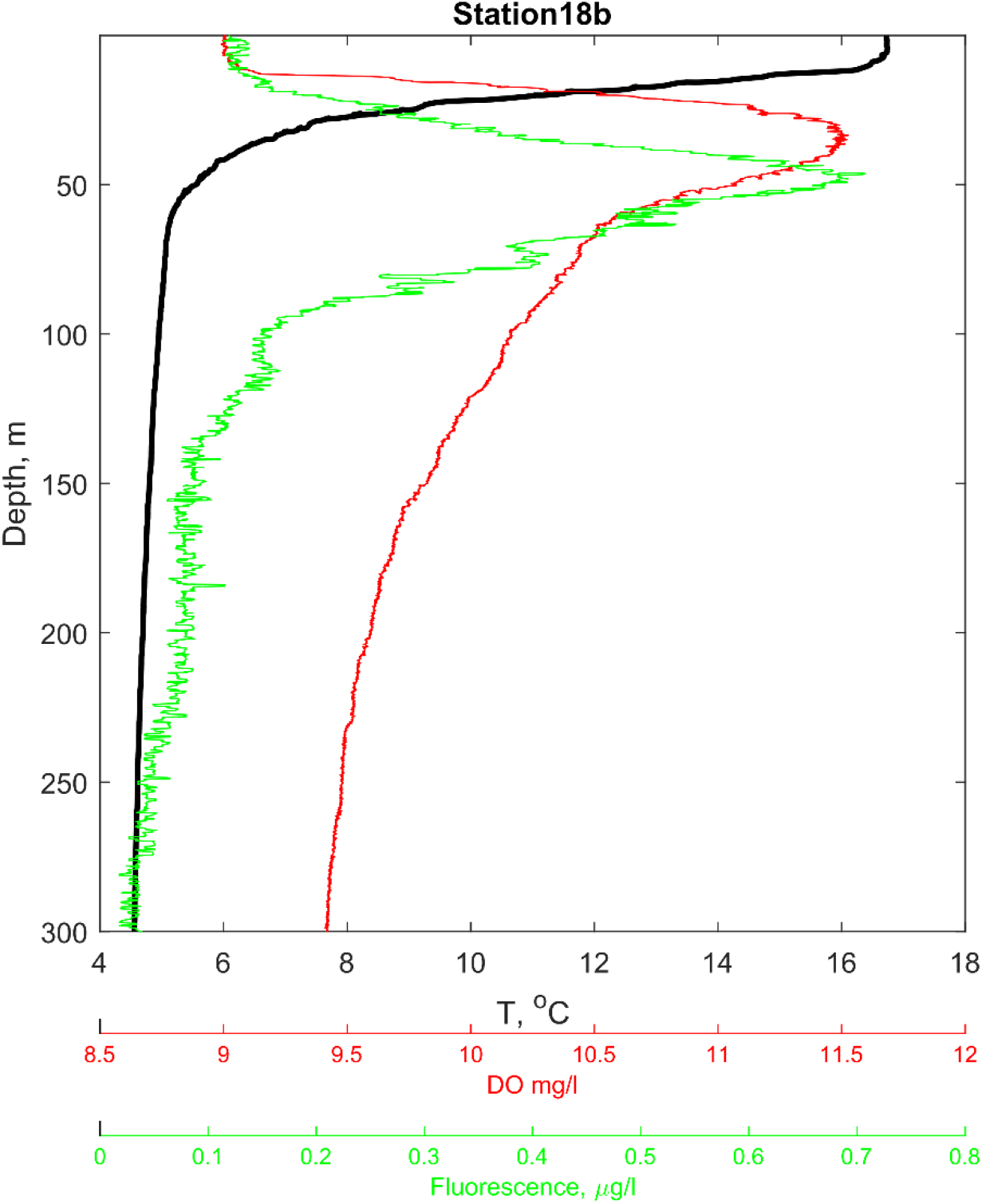
Temperature (black), dissolved oxygen (red), and fluorescence profiles in the upper 300 m of the central part of Issyk-Kul (18b in Fig. 1).

